# Notch dimerization provides robustness against environmental insults and is required for vascular integrity

**DOI:** 10.1101/2024.04.26.591315

**Authors:** Kristina Preusse, Kim Cochran, Quanhui Dai, Raphael Kopan

**Affiliations:** Division of Developmental Biology, Department of Pediatrics, University of Cincinnati College of Medicine and Cincinnati Children’s Hospital Medical Center, Cincinnati, Ohio, USA; State Key Laboratory of Genetic Engineering, School of Life Sciences, Greater Bay Area Institute of Precision Medicine (Guangzhou), Zhongshan Hospital, Fudan University, Shanghai 200438, China

## Abstract

The Notch intracellular domain (NICD) regulates gene expression during development and homeostasis in a transcription factor complex that binds DNA either as monomer, or cooperatively as dimers. Mice expressing <u>N</u>otch <u>d</u>imerization-<u>d</u>eficient (NDD) alleles of Notch1 and Notch2 have defects in multiple tissues that are sensitized to environmental insults. Here, we report that cardiac phenotypes and DSS (Dextran Sodium Sulfate) sensitivity in NDD mice can be ameliorated by housing mice under hypo-allergenic conditions (food/bedding). However, compound heterozygote NDD mice (N1^RA/–^; N2^RA/–^) in hypo-allergenic conditions subsequently develop severe hydrocephalus and hemorrhages. Further analysis revealed multiple vascular phenotypes in NDD mice including leakage, malformations of brain vasculature, and vasodilation in kidneys, leading to demise around P21. This mouse model is thus a hypomorphic allele useful to analyze vascular phenotypes and gene-environment interactions. The possibility of a non-canonical Notch signal regulating barrier formation in the gut, skin, and blood systems is discussed.

## INTRODUCTION

The Notch signaling pathway controls a wide variety of developmental and homeostatic processes via ligand induced proteolysis of the Notch receptor. Mammals encode four receptors (Notch1 (N1), Notch2 (N2), Notch3 (N3) and Notch4 (N4)) and five ligands (Delta like1 (Dll1), Delta like3 (Dll3), Delta like4 (Dll4), Jagged1 (Jag1) and Jagged2 (Jag2)). Receptors and ligands are Type I transmembrane proteins that mediate signal transduction between adjacent cells, triggered by endocytosis of a Ligand-bound receptor and subsequent conformational change in the Notch LNR/NRR (Lin12-Notch-repeat/Notch negative regulatory region) domain. This conformation change enables proteolytic cleavage of Notch mediated by ADAM10 (ADAM metallopeptidase domain 10). Subsequently, the γ-secretase complex cleaves the transmembrane domain to release the Notch intracellular domain (NICD), which translocates to the nucleus where it interacts with the DNA binding protein RBPjK (Recombinant binding protein for immunoglobulin Kappa j region, which is also known as CSL (CBF1/Suppressor of Hairless/LAG-1)) and co-activators (Mastermind-like (MAML) family members) to assemble the Notch transcription complex (NTC). The NTC binds enhancers and promoters to activate transcription of target genes [1–6]. NTCs can activate transcription via two type of DNA sites: independent CSL sites recruit monomeric NTCs, whereas two sites arranged in a head-to-head orientation 15-18 nucleotides apart (sequence-paired-sites or SPS) [7–9] recruit cooperative NTC dimers through a conserved Arginine (ARG) in the NICD ankyrin (ANK) domain [9, 10]. Cooperative binding by *Drosophila* Notch dimers on SPS endows them with improved resistance to negative regulators [11].

To understand the role of Notch dimerization in mammals, we generated mice with ARG-ALA substitution in the *Notch1* (N1^R1974A^, henceforth N1^RA^) and in *Notch2* (N2^R1934A^, or N2^RA^) genes, preventing homo- and hetero-dimerization [9, 12]. The intracellular domains (ICDs) can form NTCs, leaving Notch functionality at CSL sites unaffected [13], but abolishing cooperative binding on SPSs. Mice homozygous for <u>N</u>otch <u>d</u>imerization-<u>d</u>eficient alleles (NDD) for both N1 and N2 (N1^RA/RA^; N2^RA/RA^) are viable and fertile with a normal lifespan. However, they display enhanced sensitivity to environmental irritants: fur mite infestation triggered MZB expansion, seen also after exposure to house dust mite (HDM) extract. NDD mice also exhibit hypersensitivity to DSS-induced colitis and are unable to survive prolonged exposure to 1% DSS. Where compound heterozygote mice (N1^+/–^; N2^+/–^) thrive, compound heterozygote NDD mice (N1^RA/–^; N2^RA/–^) develop to term but perish postnatally from ventricular septal defects (VSD) and cardiac insufficiency [13]. In addition, while intestinal development in utero appears normal, the few surviving pups show decreased intestinal and colonic stem cell numbers [13], the molecular basis of which is due to elevated innate immune response to symbionts as described in the accompanying manuscript (Dai et. al).

The well-being of an organism is influenced by the interplay of their genome and the environment. Environmental risk factors known to impact inflammatory bowel disease (IBD) include diet, antibiotics, and NSAIDs. Diet can affect disease development and progression in IBD patients [14, 15], as evident from varying prevalence between countries [16, 17]. Studies in rodents have confirmed that diet composition can influence colitis development, with an increased susceptibility in high fat (Western), high protein, or high sugar diets [18–24]. Food allergens can also induce colitis in sensitized mice, as shown in an OVA/CT (Ovalbumin/Cholera toxin)-based food allergy model system [25]. Additionally, food-allergy-specific IgE has been found in a subset of IBD patients [26].

In this study, we use our NDD mouse model to analyze gene-environment interactions. We show that the tolerance to DSS as well as incidence and severity of VSD can improve to wild type levels by modifying both bedding and diet, demonstrating a profound impact of experimental factors on genetic predisposition. However, while controlling VSD with bedding and diet can improve survival of N1^RA/-^; N2^RA/-^ mice, these mice develop a new pathology, namely hydrocephalus leading to lethality. Further detailed analysis of NDD mice revealed increased global permeability of blood vessels, vascular malformations in the brain and vasodilation in the kidney. Together, these results demonstrate the importance of Notch dimerization in maintaining barrier integrity in the gut, in vascular networks, and in homeostasis of the postnatal brain.

## RESULTS

### Environment influences DSS sensitivity in Notch dimer deficient (NDD) mice

Notch dimerization deficiency results in enhanced sensitivity to environmental and genetic insults such as mite infestation, DSS treatment, or gene dosage. NDD mice are prone to developing colitis with severe inflammation and weight loss when given 1% DSS, a pathology wild type control mice develop only after exposure to 2.5 % DSS (previously reported in [13] and Fig 1A). To determine if modifying diet and bedding alters sensitivity to DSS, we co-housed wild type and N1^RA/RA^; N2^RA/RA^ mice under three conditions: 1) autoclaved diet and corn cob bedding (standard housing), 2) autoclaved diet and iso-pad bedding, and 3) extruded diet and iso-pad bedding (hypo-allergenic conditions). Both diets used are considered standard rodent chow, but the extruded diet lacks fish meal and soy products (S1 Table), which can be a source of the pro-inflammatory molecules nitrosamines and isoflavones [27–29]. Genetic background and sex influences barrier formation, DSS sensitivity, microbiome composition and cytokine expression [30–33]; therefore, to reduce their confounding effects, we used isogenic C57BL/6 male mice for all DSS treatment analyses unless otherwise stated. Mice were housed under each respective housing condition for a minimum of 4 weeks before treatment with 1% or 2.5% DSS.

**Fig1.**
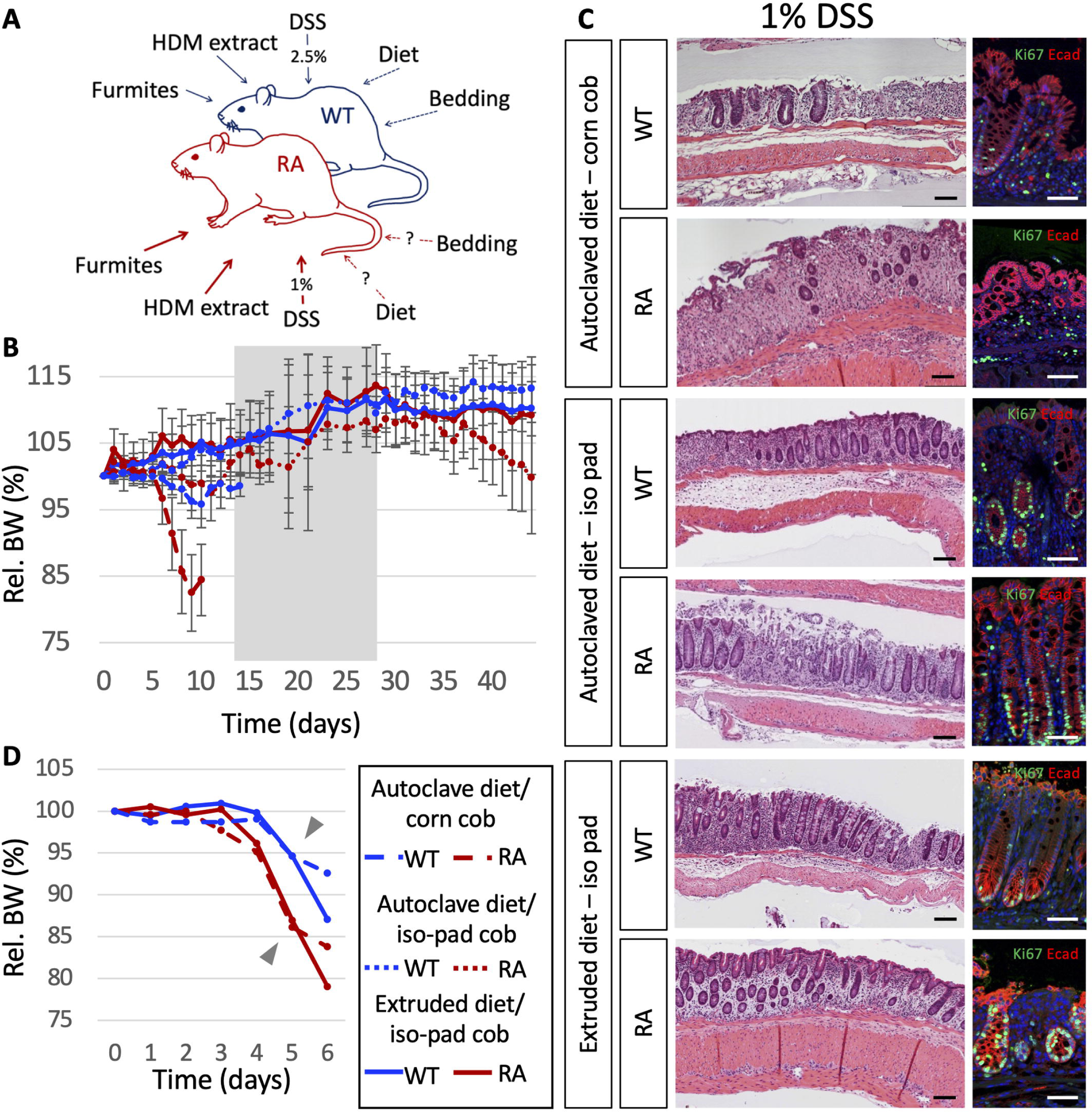
Notch dimer-deficient mice are sensitive to environment. **A**. Notch dimer-deficient mice (RA) are hypersensitive to fur mite infestation (or HDM treatment) and to DSS as reflected by the arrow weight). Development of colitis in wild type mice is evident after exposure to 2.5 % DSS, RA cannot tolerate exposure to 1 % DSS. The effect of diet and bedding, two environmental factors affecting humans, is analyzed in this study. **B**. Percent body weight relative to starting weight at d0 of mice treated with 1 % DSS for 14 days, rested with normal drinking for 14 days (gray) and subjected to a second round of 1 % DSS for 14 days. **C**. Histological analysis by H&E stain (scale bars 100 μm) and immunofluorescence (scale bars 50 μM) for E-Cadherin (red), Ki67 (green) and DAPI (blue) of distal colon sections from mice after two rounds of 1 % DSS. **D**. WT and RA mice housed with iso-pad bedding and extruded diet were treated with 2.5% DSS for 7 days, bodyweight relative to start day is shown. Abbreviations: BW: body weight, RA: N1^RA/RA^; N2^RA/RA^, WT: wild type, DSS: dextran sodium sulfate, HDM: house dust mite.

NDD mice in standard housing given 1% DSS in drinking water start losing weight after 6 days and had to be euthanized after 7 to 9 days due to severe weight loss and blood in feces, indicators of severe colitis (Fig 1B, 1C). Co-housed wild type mice show only minor weight loss during 14 days of 1% DSS treatment. On iso-pad bedding, control wild type mice showed no ill effects to 1% DSS regardless of diet (Fig 1B). When housed on iso-pad bedding, fed with autoclaved diet, and given 1% DSS in drinking water N1^RA/RA^; N2^RA/RA^ mice lose weight at a slower rate than on corn cob bedding, noticeable around day 8 of 1% DSS, (Fig 1B, 1C) and recover when given DSS-free water for 14 days (gray zone in Fig. 1B). A second 14-day treatment with 1% DSS on autoclaved chow and iso-pad bedding leads to weight loss in N1^RA/RA^; N2^RA/RA^ mice after 7 to 8 days. By contrast, NDD mice in hypoallergenic conditions are minimally affected by 1% DSS (Fig 1B). Histological analyses confirm the presence of inflammation in the colon in all DSS treated mice (Fig 1C). Immunofluorescence detection of Ki67 (green, Fig 1C) in mice housed in standard conditions showed fewer proliferation-competent stem cells in N1^RA/RA^; N2^RA/RA^ mice compared to wild type mice after one round of 1% DSS. By contrast, even after two rounds of 1% DSS in mice housed on iso-pad bedding, regardless of diet, the fraction of Ki67 positive cells in the NDD crypts is comparable to controls (Fig 1C). Notably, NDD mice kept in hypoallergenic conditions are still more sensitive to 2.5% DSS than co-housed wild type mice (Fig. 1D), highlighting their genetic predisposition. This data confirms a sensitization to DSS in NDD mice and shows that the environment (bedding and chow) can have profound effects on experimental outcomes: hypo-allergenic bedding and diet modulate the severity of DSS sensitivity inherent to NDD mice. Stated differently, NDD mice are hyper-sensitive to their environment.

### Environment and Notch-dimerization influence cytokine profiles in mice

To determine if diet influenced environmental sensitivity in NDD mice via humeral factors, we compared serum cytokine levels in NDD and wild type mice kept in standard housing, to the serum levels of NDD and wild type mice housed in hypo-allergenic conditions. We tested serum collected from dams at E16.5 to account for factors that could influence offspring during development and lactation. Overall, the changes in cytokine levels due to genotype and environment are small (S3 Fig). Cytokines altered by environmental conditions could be grouped into two classes: diet-dependent, induced independent of genotype (S3 Fig A) and genotype-dependent, affecting all independent of diet and bedding (S3 Fig B). Mice fed with extruded diet showed significantly enhanced Amphiregulin (AREG), Fibroblast growth factor 21 (FGF-21) and Leptin (Lep) levels in the blood (S3 Fig A). Regenerating islet-derived protein 3 gamma (Reg3g) and Osteopontin (Opn) levels, which are associated with senescence [34], were elevated in NDD compared to wild type mice (S3 Fig B). These findings reinforce the conclusions of the accompanying paper (Dai et al.), namely, that cell autonomous elevation of the innate immune defense apparatus within intestinal epithelial cells is likely to confers hypersensitivity.

### Changes in diet and bedding improve survival of N1^RA/–^; N2^RA/–^ mice, exposing propensity towards the development of hydrocephalous

We previously reported that hemizygous N1^RA/–^; N2^RA/–^ mice on a mixed background die at P1 due to severe VSDs, and the few escaping VSDs die due to complete disintegration of their GI (gastrointestinal) tract [13]. To test if the hypo-allergenic environment also reduces the incidence of VSD and improves survival of N1^RA/–^; N2^RA/–^ mice, we crossed N1^RA/RA^; N2^RA/RA^ mice with N1^+/–^; N2^+/–^ housed on iso-pad bedding and fed the extruded diet. We calculated the ratio of expected genotypes among surviving pups on a mixed genetic background at P1 (as done in [13]). N1^+/–^; N2^+/–^ controls were born at the expected ratio (Chi^2^ p=0.74, S2 Table B). Whereas previously only 2/14 N1^RA/–^; N2^RA/–^ pups were found alive at P1 [13], now 16/35 survive up to postnatal day 18 to 21 (P18-P21; Fig 2). N1^RA/–^; N2^RA/–^ pups found dead at P1 had VSD (Fig 2B) as we reported previously [13]. The number of surviving N1^RA/–^; N2^RA/–^ pups was still significantly lower than expected (S2 Tab A), but much better than previously reported in standard housing conditions [13] (p=10^−5^ compared to p=10^−9^; comparison Chi^2^ p=10^−11^, S2 Tab C). Surviving N1^RA/–^; N2^RA/–^ mice were smaller than littermates at P18 (Fig 2A); we analyzed hearts and intestines of hemizygotes N1^RA/–^; N2^RA/–^ and N1^RA/–^; N2^+/RA^ to ask if poor growth could be attributed to other GI or heart defects. Histological analysis of the intestine did not show any defects in hemizygotes when kept in hypo-allergenic housing, confirming rescue of the dosage effect on the GI track (Fig. 2C). Hearts of surviving N1^RA/–^; N2^RA/–^ pups were histologically normal (Fig. 2B), however, by the third week of life they displayed dome shaped heads and had to be euthanized due to poor health (Fig. 2D). Coronal sections through the brains identified severely enlarged ventricles and hemorrhages in the brain cortex (Fig 2E). Taken together, our data shows that hypoallergenic housing and changes in diet resulted in a partial rescue of heart and gut development of N1^RA/–^; N2^RA/–^ mice with improved survival. However, these mice eventually succumb to postnatal hydrocephalous.

**Fig2.**
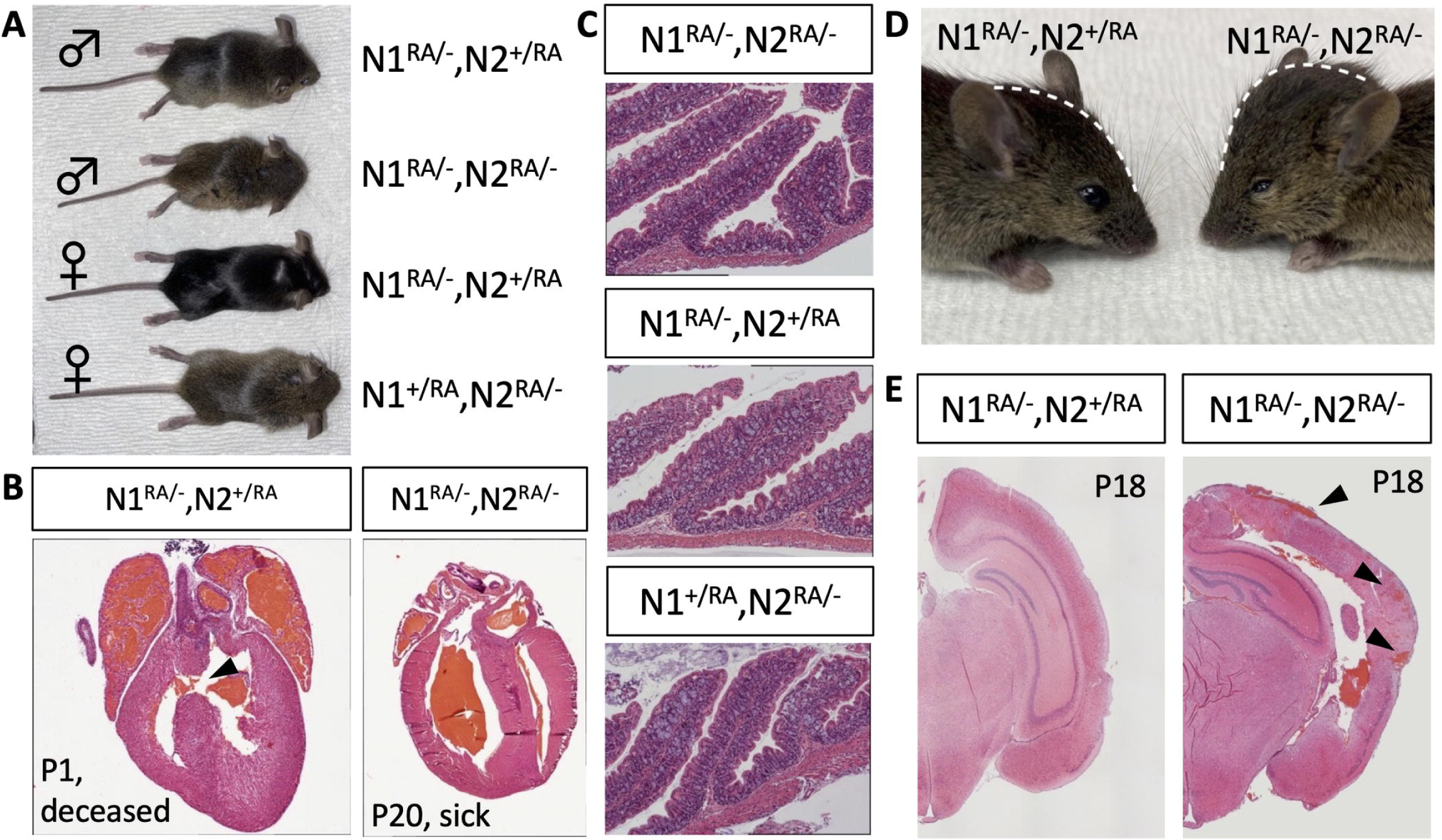
Improved survival of N1^RA/-^, N2^RA/-^ mice on anti-inflammatory diet and bedding exposes a hydrocephalous phenotype. **A**. A subset of hemizygous N1^RA/-^; N2^RA/-^ mice survive to P18 but are noticeably smaller than littermates. **B**. Hearts of surviving pups appear normal, but N1^RA/-^; N2^RA/-^ litter mates that die at P1 or P2 show ventricle septal defects (arrowhead). **C**. The gut morphology of surviving P10 N1^RA/-^; N2^RA/-^ mice appears normal in H&E stains. **D**. Heads of surviving N1^RA/-^; N2^RA/-^ mice are dome shaped at P18. **E**. H&E stains of coronal brain sections show enlarged cavities in brain ventricles with hemorrhagic brain tissue (arrowheads) not detected in litter mate controls.

### Dimer deficiency causes vascular defects in brain and kidney

To analyze if vascular leakage contributed to hydrocephalus, blood vessel integrity in ten N1^RA/RA^; N2^RA/RA^ mice and five unrelated but co-housed wild type controls on the C57BL/6 background was evaluated by trans-cardiac perfusion with Evans blue [35, 36]. No leakage was observed in pre-cleared or cleared tissue from controls, and whole mount brain imaging showed that the blue stain was confined to blood vessels (Fig 3A). By contrast, a subset (4/10) of perfused NDD mice showed evidence of vascular leakage in brain tissue pre-clearing (Fig 3A) with leakage detected in 8/10 after clearing. The severity of vascular leakage varied in NDD mice from a few small blue areas to completely blue brains (Fig 3B, total 8/10 p=0.007 Fisher’s exact test), consistent with a compromised blood brain barrier (BBB). Similarly, no leakage was observed in perfused and cleared control kidneys, whereas 50% of NDD (3/6) kidneys showed evidence of leakage (Fig 3C). We next looked for leakage after trans-cardiac perfusion with Evans blue in mice in which a null allele replaced either the wild type *Notch1* or *Notch2* allele. Uncleared perfused kidneys of N1^+/RA^; N2^RA/–^ mice resembled wild type controls (Fig 3D), and after clearing, vascular leakage was detected in only 2 out of 8 animals (Fig 3E, 6/8 without leakage). By contrast, blue background stain can be observed in most (5/8, p=0.07 Fisher’s exact test) kidneys isolated from N1^RA/-^; N2^+/RA^ (Fig 3D, 3E). These data suggest that *Notch1* dimerization and dosage in endothelial cells is more critical than *Notch2* dosage in vSMCs, but both cell types cooperate to promote vascular integrity in multiple organs.

**Fig3.**
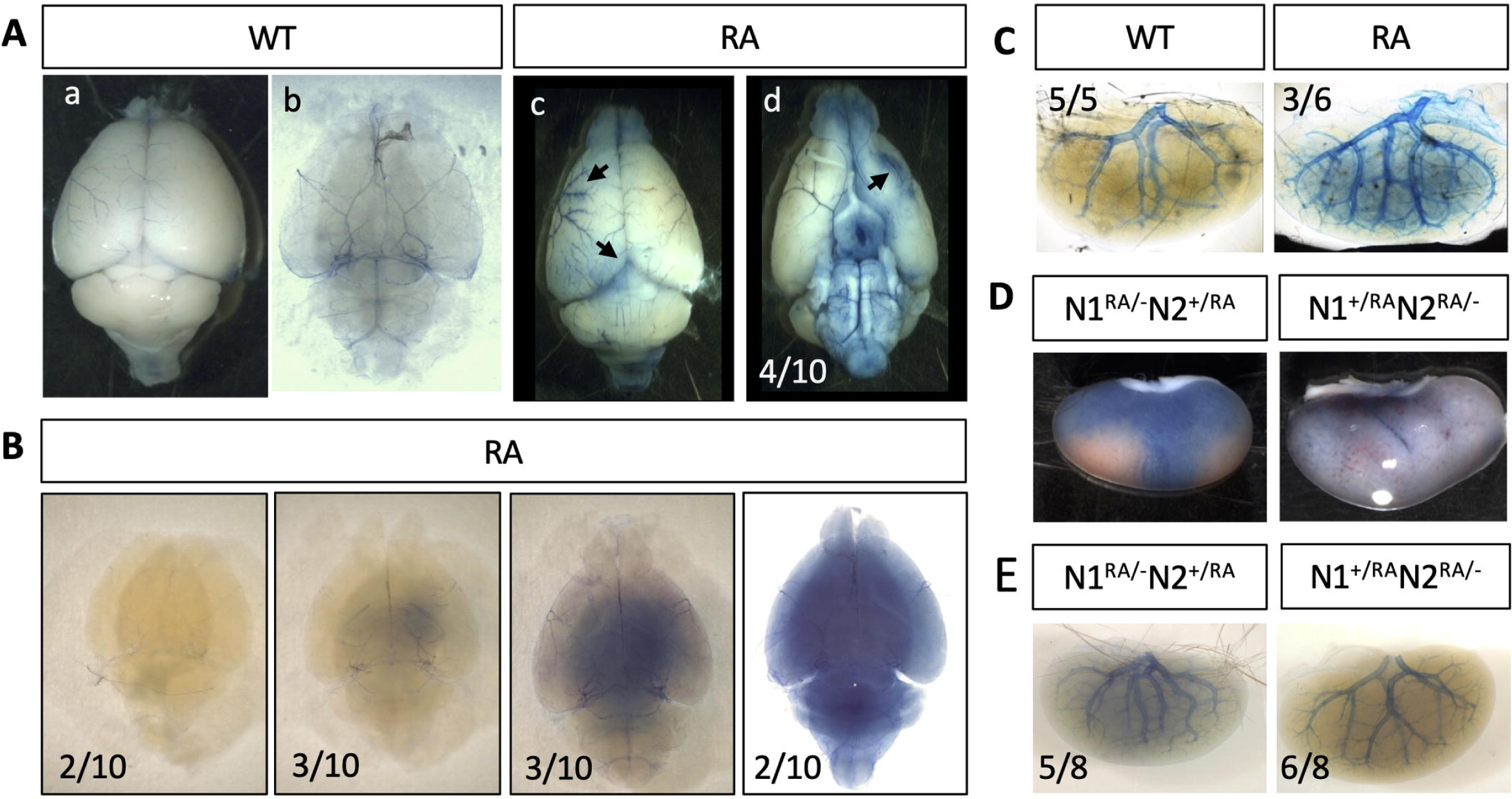
Deficiency in Notch1 dimerization causes vascular leakage. **A**. Evans blue perfusion labels blood vessels in brains of control and NDD mice. **B**. Cleared brains from Evans blue perfused NDD mice display various degrees of vascular leakage (total 8/10, p=0.007 – Fisher’s exact test). **C**. Kidneys from Evans blue perfused mice (5/5 vs 3/6 p=0.1818, Fisher’s exact test). **D**. Kidneys from N1^RA/-^; N2^+/RA^ mice display blue patches when perfused with Evans blue (5/8), only 2/8 of N1^+/RA^; N2^RA/lacZ^ kidneys show leakage. **E**. Vascular leakage is detected in subset of N1^RA/-^; N2^+/RA^ kidneys, most N1^+/RA^; N2^RA/lacZ^ look comparable to wild type.

Analysis of cerebral vasculature identified disruption of the Circle of Willis (CoW) patterning when Notch signaling was inactivated in vascular smooth muscle cells (vSMCs) by conditional overexpression of dominant negative Mastermind (dnMAML) [37]. To test if Notch dimerization in ECs (*Notch1*) and vSMC (predominantly *Notch2*) contributed to vascular morphology in the CoW, we examined Evans blue perfused brains (Fig 4A, 4B). The anterior part of the CoW contains the anterior cerebral arteries (ACA, arrows) and anterior communicating arteries (AcoA, arrowheads) (Fig 4Ba). Control brains showed mostly symmetric arrangement of the major blood vessels (Fig 4Ba). By contrast, N1^RA/RA^; N2^RA/RA^ mice displayed a range of vascular morphologies, from small irregularities that were otherwise like wild type, to very irregular and/or asymmetric vasculature (Fig 4A, 4B). The degree of malformation in N1^RA/RA^; N2^RA/RA^ mice varies between asymmetric differences in vascular diameter (Fig. 4Bb, asterisk), to fusion between different arteries (Fig 4Bc, asterisk), to completely asymmetric vessel arrangement (Fig. 4Bd, asterisk). Reduction in either N1 or N2 (N1^RA/–^; N2^+/RA^ and N1^+/RA^; N2^RA/–^, respectively), can cause vascular malformations in the CoW with about 50% penetrance (Fig 4Be and 4Bf), consistent with a role for each receptor in regulating vascular patterning.

**Fig4.**
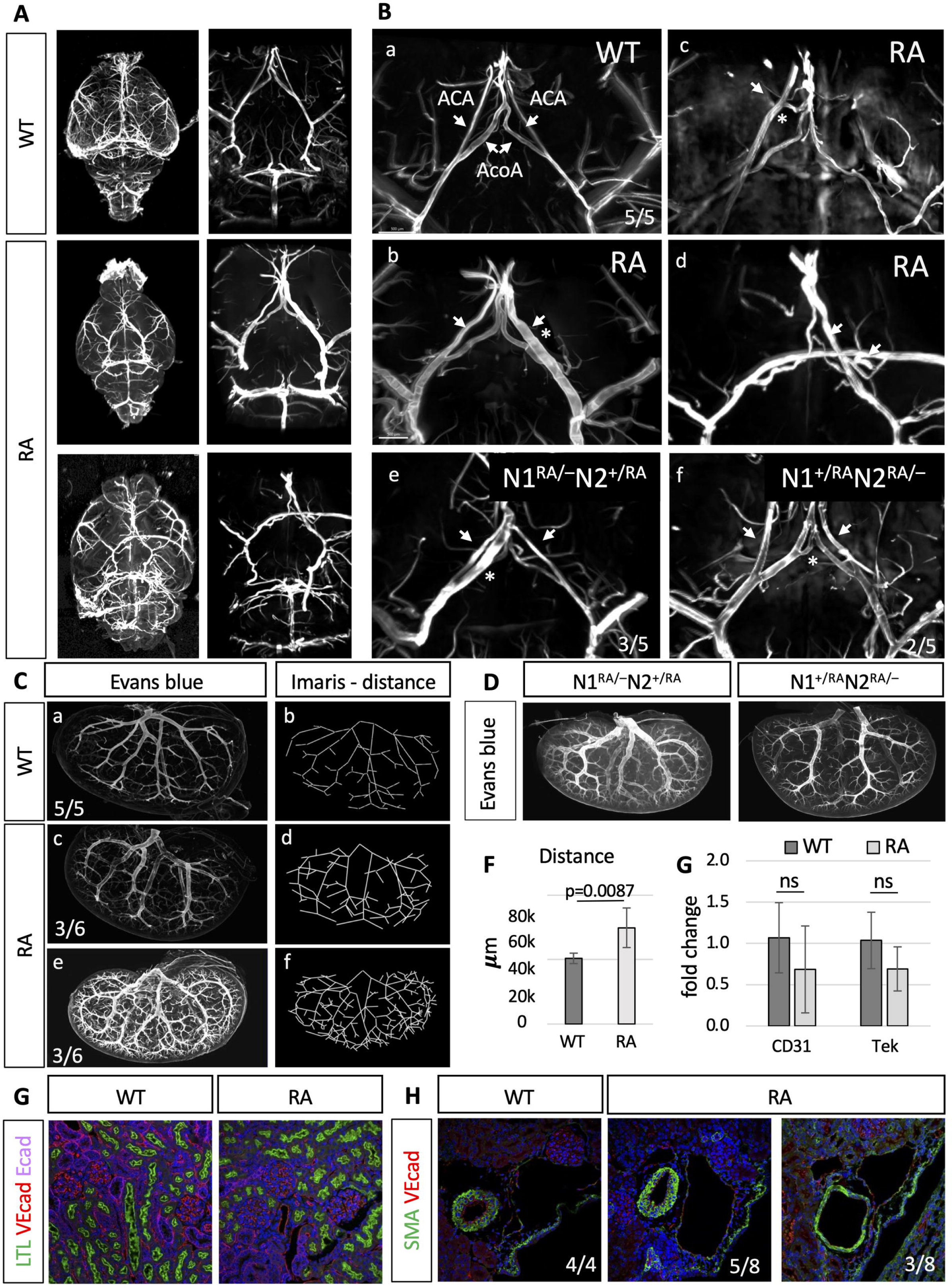
Notch dimer deficient mice display smooth muscle cell associated phenotypes in brain and kidney. **A**. Evans blue labeled vasculature in brains of dimer deficient mice show a range of irregularities compared to wild type; Circle of Willis (CoW) shown in higher magnification. **B**. Anterior part of the CoW in (a) wild type (n=5) and (b-d) dimer deficient mice (n=6). Dimer deficient mice show various degrees of malformations: (b) unilateral increased vessel thickness of the ACA, (c) fusion between ACA and AcoA and (d) complete asymmetry; arrows indicate ACA, stars indicate malformations (p=0.022, Fisher’s exact test) **C**. Evans blue labeled vasculature of kidneys from wt and RA mice (a,c,d) and vascular branching displayed using Imaris measuring tool (b,d,f). Dimer mutant RA mice display increased labeling with 50% penetrance (e,f). **D**. Evans blue labeled vasculature in N1^RA/-^; N2^+/RA^ or N1^+/RA^; N2^RA/lacZ^ kidneys. **E**. Distance measurements of Evans blue stained vasculature in kidneys shows increase in dimer mutants compared to wild type. **F**. qPCR analysis for endothelial markers in WT and RA kidneys. **G**. Immunofluorescence staining of sections through the cortex of WT and RA kidneys vascular endothelial cells (VE cadherin, red, LTL, green and, ECadherin in blue). **H**. Sections of kidney blood vessels show vasodilation in a subset of dimer deficient mice. Abbreviations: ACA: Anterior cerebral artery, AcoA: Anterior communicating artery, CoW: Circle of Willis, RA: N1^RA/RA^; N2^RA/RA^, WT: wild type.

As was reported for mice with a defect in SMCs [38–40], hydronephrosis was frequently seen in NDD mice (S4 Fig A and B). Together, the vascular malformations in the CoW and the development of hydronephrosis suggest a defect in SMCs in NDD mice. To ask if SMCs are affected, we performed histological analyses of kidney sections to examine the SMC-rich ureter. Branching morphogenesis and glomerular numbers were the same whether N1^RA/RA^; N2^RA/RA^ mice had hydronephrosis or not (S4 Fig C). Cross sections of the ureters uncovered no morphological difference that segregated with the RA alleles or with hydronephrosis. Likewise, similar numbers of SMCs were found in ureter cross sections stained by immunofluorescence for smooth muscle actin (SMA) and transgelin (Tagln) (S4 Fig D).

Similar to CoW malformation, we observe denser kidney vasculature in 50% of the Evans blue perfused RA mice (Fig 4C). Labeled blood vessels in perfused kidneys from N1^RA/–^; N2^+/RA^ or N1^+/RA^; N2^RA/–^ mice do not show increased Evans blue stained blood vessel numbers (Fig 4D). To quantify the visible vasculature, we used the Imaris (Oxford Instruments) measurement tool. Distance measurements confirmed an elevated label density in a subset of analyzed N1^RA/RA^; N2^RA/RA^ mice in comparison to wild type controls, reaching deeper into smaller diameter vessels (Fig. 4 Cb,d,f and Fig 4E). To ask if the increased vessel density reflected a change in EC numbers and therefore the amount of blood vessels, we performed quantitative RT-PCR analysis for two endothelial markers, the platelet and endothelial cell adhesion molecule 1 (PECAM1 or CD31) and TEK receptor tyrosine kinase (Tek). This analysis suggested that the number of ECs was not increased in N1^RA/RA^; N2^RA/RA^ mice (Fig 4F). Similarly, immunofluorescent staining of kidney sections for Vascular endothelial cadherin (VEcad) in kidneys from N1^RA/RA^; N2^RA/RA^ and control mice were indistinguishable (Fig 4G). The increase in label density without an overall increase in blood vessel numbers suggests perfusion in RA mice is more complete compared to controls, perhaps due to altered vascular tone. Cross sections through larger kidney vessels stained for VE cadherin (VEcad) and Smooth muscle actin (SMA) identified dilation of blood vessels in some NDD mice (Fig 4H), consistent with improved perfusion due to vasodilation.

## DISCUSSION

### The environment modulates homeostasis of multiple organs in dimer-deficient mice

In a previous study, we described a mouse model with point mutations in N1 and N2, preventing homo- or heterodimerization [13]. When not challenged, N1^RA/RA^; N2^RA/RA^ (NDD) animals are viable and show no overt signs of impaired health. Developmental decisions *in utero* are robust and indifferent to Notch dimerization. We have shown that postnatally, NDD mice are hyper-sensitive to environmental insults with variable, organ specific severity. When exposed to fur mites or HDM extract, NDD mice show expansion of MZB cells, a phenotype associated with Notch gain of function; chronic exposure to fur mites obliterates spleen architecture by massive expansion of MZB cells [13]. A reduction of stem cells was observed in the colon, and exposure to low doses of DSS triggers severe colitis [13], a phenotype compatible with Notch loss of function [41–43]. To analyze the effect of environmental conditions on NDD mice, we systematically tested the effect of hypo-allergenic diet and bedding (S1 Tab). While the rodent housing conditions compared in this study are all considered conventional in animal husbandry, the change to hypo-allergenic conditions significantly improved tolerance to DSS in homozygous NDD mice (Fig 1), as well as lowered the frequency of VSDs, and improved survival of compound heterozygote N1^RA/-^; N2^RA/-^ animals (Fig 2), demonstrating how important animal housing is for experimental outcomes (and by extension, to human health). Isolated gut epithelial spheroids from NDD mice show elevated innate immune defense against symbionts (accompanying manuscript – Dai et al.) which explains the exacerbated response to mucosal stripping. In addition, hypo-allergenic housing conditions induced tolerance to DSS and lowered VSD frequency may also be explained by elevated levels of Amphiregulin (AREG) in NDD mice fed with extruded (hypo-allergenic) diet (S3 Fig A), consistent with findings that experimental increase of AREG levels can rescue DSS-induced colitis in mice [44] and improve intestinal regeneration [45]. Correspondingly, deletion or reduction of the murine AREG-EGFR signaling axis increased susceptibility to DSS [46, 47].

### Notch dimerization is a modifier of vascular integrity

Unlike N1^RA/RA^; N2^RA/RA^ or N1^+/–^; N2^+/–^mice, N1^RA/-^; N2^RA/-^ pups fail to survive postnatally [13]. The hypo-allergenic housing conditions improved survival of N1^RA/-^; N2^RA/-^ mice leading to the discovery of vascular phenotypes, namely, hydrocephalus (Fig 2) and more generally, loss of endothelial barrier function in homozygous NDD mice (Fig 3 and 4). Retrospective analysis revealed that vascular defects were also present in N1^RA/RA^; N2^RA/RA^ mice (Fig. 3 and 4), though they did not compromise viability. Vascular defects occur with roughly 50% penetrance and include vascular leakage, malformations in the Circle of Willis (CoW) and vasodilation in the kidney (comprehensive analyses may reveal additional pathologies). The defects are related to reduced Notch1 function in ECs [48, 49] and augmented by reduced Notch2 function in vSMCs [50–52], with carriers of RA substitutions in both N1 and N2 being the most severely affected.

Impaired vascular barrier function in NDD mice is due to loss of Notch activity. N1 signaling is activated in ECs by sheer stress and has been shown to lower vascular permeability through improvement of cell-cell-junctions [53, 54]. Non-canonical Notch signaling [53] (see below) as well as Calcium regulation [54] were proposed to be involved in executing these effects. Similarly, in *in vitro* experiments, Dll4-Notch signaling in ECs was shown to promote barrier function through induction of VE cadherin expression [55]. By contrast, exogenous ligand-mediated Notch activation *in vitro* increases permeability of endothelial monolayers *in vitro* [56], suggesting Notch signal strength, perhaps both transcriptional (canonical) and non-canonical, modulates vascular permeability.

Apart from vascular leakage, we observed vascular malformations in the CoW as well as vasodilation in a subset of NDD mice (Fig 4), consistent with contributions from both ECs and vSMCs. The observed vascular malformations in the CoW resemble previously described phenotypes for pan-Notch signaling inhibition in embryos and adult mice by vSMCs-specific expression of dnMAML [37, 57], an orthogonal conformation that NDD mice are hypomorphic transcriptionally in SMCs. Similarly, loss of RBPjK in pericytes leads to formation of arterio-venous malformations [58]. Moreover, postnatal deletion of RBPjK [59, 60] or N1 [59, 61] in ECs causes defects in vasculature including enlarged vessels, shunts and abnormal vascular architecture [59, 60] as well as changes from arterial to venous vessel identity with signs of inflammation [62]. Conversely, constitutively activated Notch signaling in ECs also causes vascular defects with shunting and vessel enlargements [63], which can be reversed by later deletion of Notch signaling [64]. This underscores the importance of Notch signaling strength in vasculature, with both gain and loss of function causing malformations.

The observed increase in perfused vessels in 50% of kidneys isolated from N1^RA/RA^; N2^RA/RA^ mice without a change in vessel or ECs numbers suggested changes in vascular tone. Notch signaling in vSMCs regulates vascular tone by affecting dilation and constriction as it is required for the response of vaso-regulators and vasoactive genes [65]. Inhibition of Notch signaling upregulates the vasodilators adrenomedullin and neurotensin and downregulates the vasoconstrictor angiotensin II [65]. Inhibition or vSMC-specific expression of dnMAML attenuates response to vasodilators or vasoconstrictors in mice [66]. Similarly, N3 null mice experience dilation of blood vessels [67, 68]. In the kidney vasculature of N3 null mice, the response to vasodilators and vasoconstrictors is dampened, resulting in mice which develop severe renal and cardiac phenotypes due to high blood pressure [69]. Conditional deletion of RBPjK in ECs postnatally at birth or in adult mice also causes dilated vessels [60], demonstrating an involvement of ECs in addition to vSMCs to controlling the vascular tone.

Vascular defects, including hemorrhages and vascular leakage, have been observed in patients with Notch-associated disease. Spontaneous hemorrhages, with potentially fatal outcomes, are observed in patients with Alagille syndrome, as well as in a mouse model for Alagille syndrome. [70–72]. Patients with Adams-Oliver Syndrome (AOS) can exhibit vascular leakage in their retinas [73], and vascular malformations in human patients, mainly arterio-venous malformations, have been linked to Notch signaling. Together, the vascular defects observed in our dimer-deficient mice resemble defects observed in human disease, and our study suggests that vascular leakage in the context of hypomorphic Notch in humans may also respond to environmental condition. Uncovering environmental modulators of vascular lockage in our model could lead to strategies ameliorating disease severity.

### Non-canonical, dimer-dependent Notch signals support Barrier formation

The hyper-sensitive RA mice have defects in their skin barrier, gut barrier, and in vascular integrity; all modulated by environmental challenges (fur mites, bedding, DSS, and diet exacerbate; hygienic conditions, hypo-allergenic diet and bedding alleviate). It has not escaped our attention that barrier defects in skin and gut share not only a common requirement for Notch dimerization but are also at a corner of vertebrate Notch biology where some part of the Notch activity appears to be delivered in an RBPjK-independent manner. Discrepancies between deletion of Notch receptors (or gamma secretase components) and RBPjK deletion in the skin were previously analyzed [74], concluding that RBPjK independent Notch function regulated the barrier in skin. Similarly, in the gut, RBPjK deletion [41] and small molecule disruption of Notch/RBPjK interactions are tolerated [75], but Notch cleavage inhibition and Notch receptors deletion are not [76, 77]. Combined, these studies are consistent with a non-canonical, dimer supported but RBPjK independent activity of Notch in skin and gut barrier. Herein, we describe a dimer dependent activity of Notch in regulation of vascular integrity, but it is unclear at present if this activity requires RBPjK. An RBPj-independent non-canonical function for Notch during vascular barrier formation was described [53], however, the mechanism envisioned therein is not feasible because gamma secretase is a processive protease [78, 79], trimming the transmembrane domain after substrate ICD release until the expulsion of a short remnant into the vesicular lumen or extracellular milieu. Whether/how non-canonical Notch signaling is affected by dimerization needs to be further investigated; of note, we detected no dimerization-dependent protein-protein interaction partner other than Notch itself in a TurboID experiment performed in gut cells (Dai et. al, accompanying manuscript). The mechanism explaining the discrepancy in RBPjK and Notch requirements for gut and skin barrier, and the dependence of barrier robustness on Notch dimerization, will have to be explored in detail elsewhere.

## MATERIAL AND METHODS

### Ethics statement

All animal studies were approved by the Institutional Animal Care and Use Committee of the Cincinnati Children’s Hospital Medical Center, Ohio, USA (IACUC2018-0105, IACUC2021-0086).

### Animals

All mice were housed at the Animal Facility at Cincinnati Children’s Hospital Medical Center. Used strains and genotyping information are listed in Key resources table (Table 1).

**Table 1.**
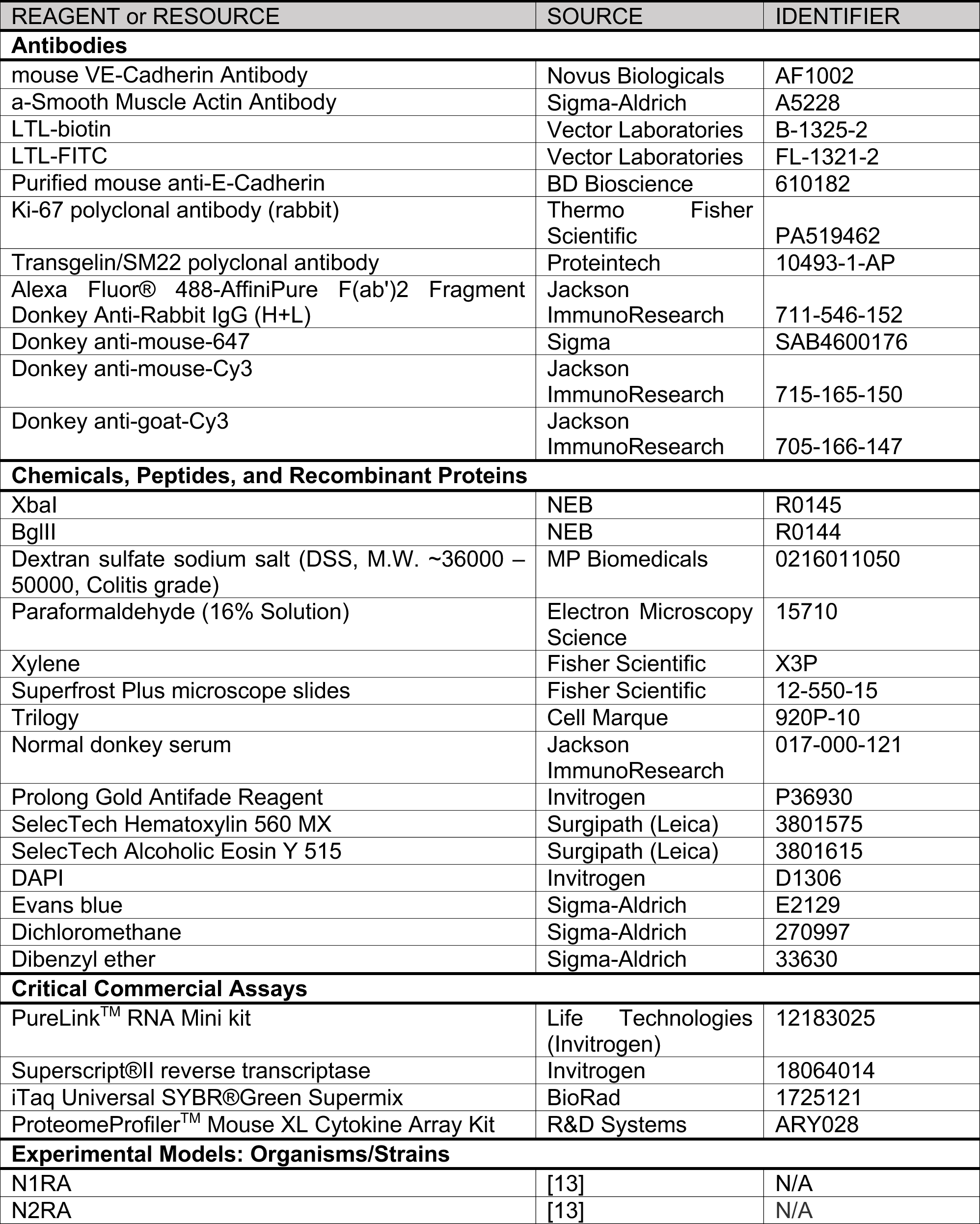
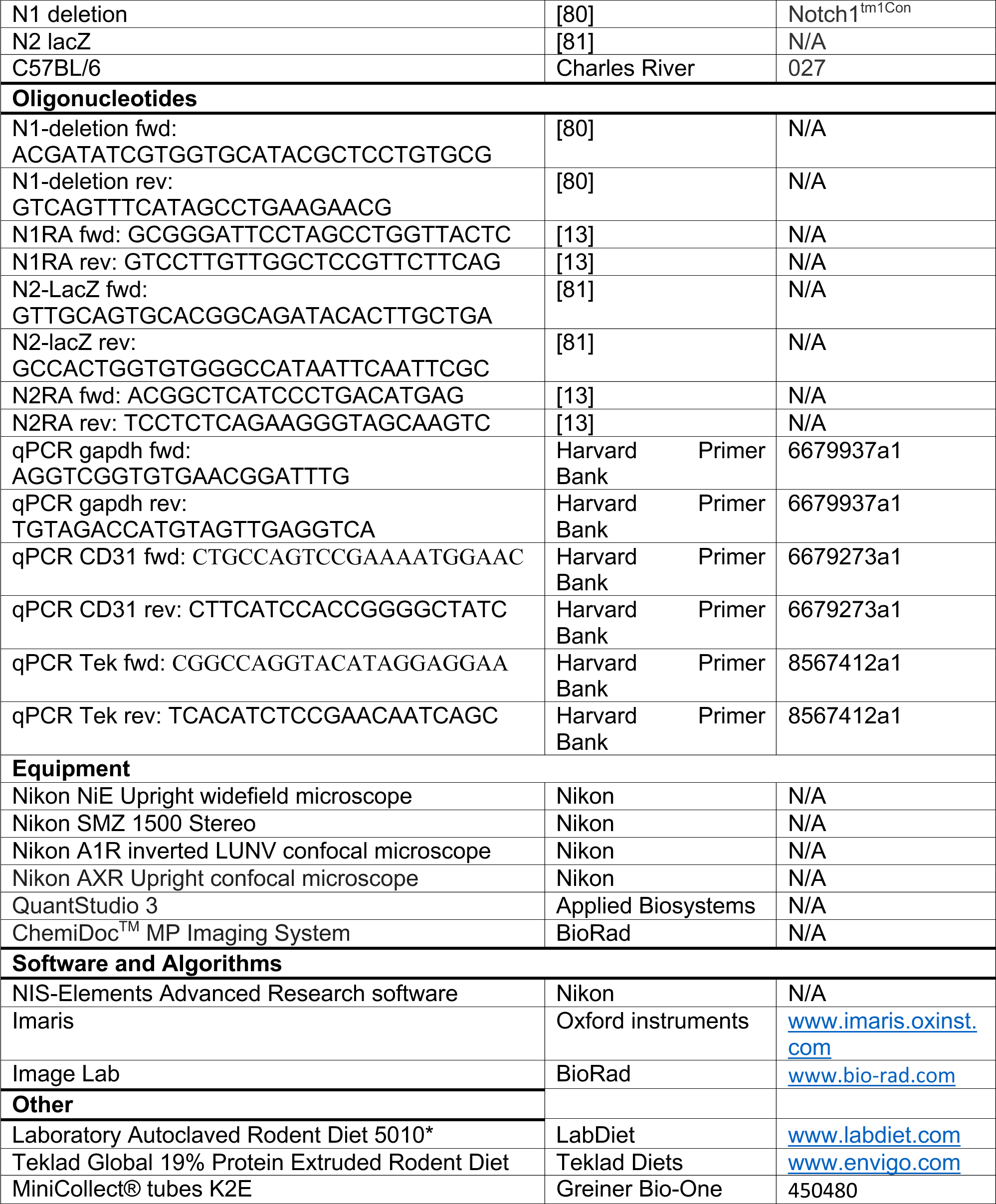
Key Resources/all used materials.

### DSS treatment

For colitis, induction mice were given 1%, 2% or 2.5% DSS (MP Biomedicals, Tab1) in autoclaved drinking water for 10 or 14 days. For long term treatment with repeated cycles of DSS, mice were given 1% DSS for 14 days, were rested with normal drinking water for 14 days and then given 1% DSS for another 14 days. If not otherwise stated, only male mice were used for DSS treatment studies. At end of treatment mice were euthanized and tissues collected for analysis.

### Tissue collection and processing

Tissue from euthanized mice was collected in ice cold PBS and fixed for 16 hours (over-night) in 4% Paraformaldehyde (PFA) at 4°C. Samples were washed with PBS and given to the Integrated Pathology Research Facility (RRID:SCR_022637) at Cincinnati Children’s for paraffin embedding and sectioning. Tissue samples for RNA extraction were flash frozen in LN2 and stored in RNA lysis buffer in −80°C until processed. For Cytokine analysis, terminal blood collection was performed from the heart, blood was collected into K2EDTA tubes (Greiner Bio-One) and pelleted for 5 min at maximum speed at room temperature. Serum samples were stored at −20°C until used.

### Evans blue perfusion

Evans blue perfusions for labeling of vasculature in mice was done as described previously [36] with small modifications. Mice of weaning age (p21) were anesthetized by intra-peritoneal injection of triple sedative solution (Ketamine/Xylazine/Acepromazine) and trans-cardiac perfused with a solution of 0.01% Evans blue (Sigma) in sterile PBS (Gibco) into the right ventricle manually using a 27G needle. Each mouse was perfused with 5-10 ml of Evans blue in PBS followed by 5 ml of 4% PFA and tissue was harvested and fixed 16h in PFA. Fixed tissue was washed in PBS for at least 24 h before tissue clearing.

### Tissue clearing (iDISCO)

Tissue clearing was done according to previously described protocol for modified iDISCO [36]. Briefly, fixed and washed tissue was dehydrated by stepwise incubation in 25%, 50%, 75% and twice in 100% methanol for 30 minutes each followed by an incubation in 50% Dichloromethane (DCM)/50% methanol for 3 hours and two times 30 min in DCM. Tissue was transferred into Dibenzyl ether (DBE) for 24-72 hours until tissue was clear and was imaged as z-stacks on a Nikon AXR upright confocal microscope with resonant scanner.

### Image processing

Images of whole organs imaged with resonant scanner were digitally de-noised and clarified using Nikon Elements software. Processed images were analyzed in Imaris. Imaris was used to display stained vasculature of the brain, and brain images were cropped in 3D to display CoW only. The Imaris surface tool was used to label stained vascular branches as well as the surface of the whole kidney. The measurement tool was used to label all stained vascular branches and measure the total length of perfused vasculature in the kidney.

### Histology

Paraffin was removed from slides with Xylene and sections were dehydrated by consecutive washes in 100%, 100%, 95%, 80% Ethanol for 2 min each. Slides were rinsed with water and stained in Hematoxylin for 4.5 min. Slides were rinsed in water, 0.05% Hydrochloric Acid, water, 0.01% Ammonium Hydroxide, water and 80% Ethanol before staining in Eosin for 20 sec. Sections were dehydrated by washing in 95% and trice in 100% Ethanol and Xylene and mounted. Stained slides were imaged on Nikon TiE widefield microscope.

### Immunofluorescence

For fluorescent immunohistochemistry, paraffin sections were de-paraffinized with Xylene and stepwise rehydrated by incubation in 100%, 75%, 50% and 25% ethanol, followed by 2 washes in PBS. Slides were boiled in Trilogy for 25 min and cooled down for 25 min for antigen retrieval. After two washes in PBS, slides were washed twice in PBS-T (PBS/0.1% Triton X100) and blocked with 10% Normal Donkey Serum (NDS) in PBS-T for 1 h at room temperature (RT). Primary antibodies, all diluted 1:100 in blocking buffer, were incubated for 16 h at 4°C. Slides were washed trice for 10 min in PBS-T, blocked again for 1 h at RT and incubated 1 h at RT with secondary antibodies diluted 1:200 in blocking buffer in the dark. Slides were washed twice with PBS-T and twice with PBS for 10 min each and incubated with DAPI in PBS for 20 min. Slides were washed three times for 30 min in PBS and mounted with ProLong Gold for imaging on Nikon A1R confocal microscope. For detailed information on used antibodies and materials see Key resource table (Tab1). Slides were imaged on Nikon A1R confocal microscope.

### RNA isolation, cDNA synthesis and qPCR

For materials used, including oligos for qPCRs, see Key resources table (Tab1). RNA extraction from mouse kidneys was performed using the PureLink RNA Mini kit (Invitrogen) according to manufacturer’s instructions. cDNA synthesis was performed with 1ug RNA and SuperScript II reverse transcriptase (Invitrogen) following manufacturer’s instructions and was diluted 1:20 in water before qPCR analysis. Quantitative PCRs were performed using Universal iTaq SYBR Green Supermix (Bio-Rad) on QuantStudio3 RT PCR system (AppliedBiosystems). Data was analyzed using the delta-delta-CT method, significance was determined by student t-test.

### Cytokine array

Cytokine analysis was performed on serum of pregnant dams on E17.5 housed on corn cob and autoclaved diet or iso-pad bedding and extruded diet. Serum from 3 mice was pooled for each array, 50ul from each animal – 150ul serum per array. Arrays were performed in duplicates. Assays were performed according to manufacturer’s instructions (R&D Systems). Briefly, membranes were blocked for 1 hour at room temperature in provided array buffer, total volume of 150ul sample per membrane were diluted in array buffer and incubated with membranes over night at 4°C. Membranes were washed in provided wash buffer before incubation with detection antibody cocktail for 1 hour at room temperature. Membranes were washed in wash buffer, incubated with Streptavidin-HRP for 30 minutes followed by multiple wash steps before imaging on BioRad Chemi Imager using Chemi Reagent Mix. Signals were quantified using Image Lab Software (BioRad), normalized to internal negative control signals and cytokine levels relative to wild type on extruded diet were calculated. Significance was determined by student t-test.

## Acknowledgements

This study was made possible, in part, using the Cincinnati Children’s Integrated Research Pathology Facility [RRID:SCR_022637] and the Bio-Imaging and Analysis Facility [RRID:SCR_022628]. We specifically acknowledge the assistance of Sarah McLeod and Matt Kofron for their guidance with tissue clearing, imaging and image analysis.

