## Supplementary material for "Notch dimerization provides robustness against environmental insults and is required for vascular integrity": S1 Fig

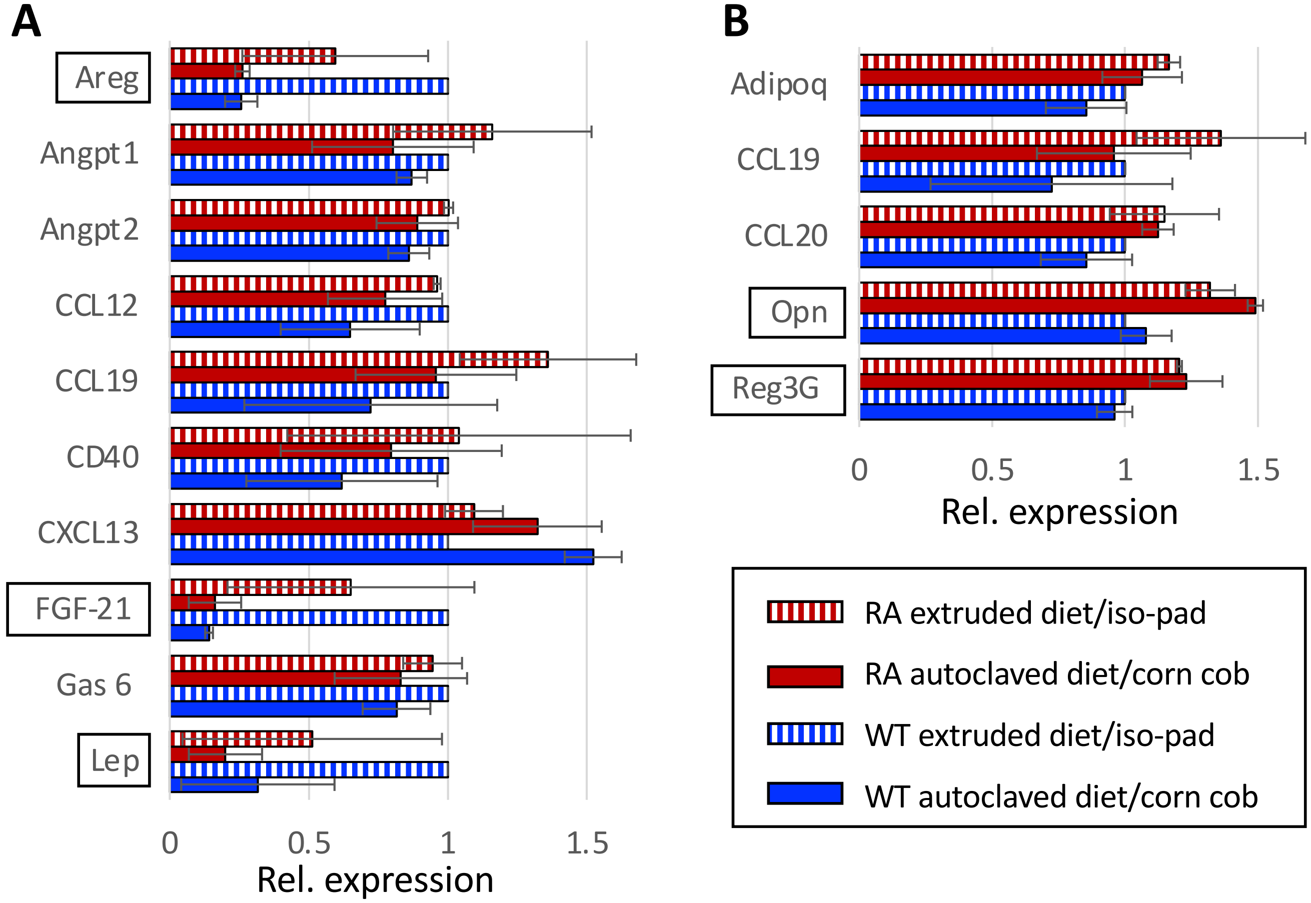


**S1 Fig. Cytokine levels in RA and WT dams are influenced by changes in diet.**

(**A, B**) Relative Cytokine expression in serum of pregnant dams housed on different bedding and fed different diet collected at E16.5/E17.5, normalized to wild type mice on extruded diet/iso-pad bedding, significant changes are indicated by boxed gene name. (**A**) Cytokines altered by diet in WT and RA mice. (**B**) Cytokines altered between WT and dimer-deficient mice independent of diet/bedding.
