## Supplementary material for "Notch dimerization provides robustness against environmental insults and is required for vascular integrity": S1 Table

**S1 Table:** Composition of autoclaved and extruded (anti-inflammatory) rodent diet, potentially inflammatory components marked in bold.

| | **Autoclaved diet** | **Extruded diet** |
|---|---|---|
| **Ingredients** | Ground corn, **dehulled soybean meal**, wheat middlings, **fish meal**, whole wheat, wheat germ, brewers dried yeast, ground oats, dehydrated alfalfa meal, **porcine animal fat** preserved with BHA and citric acid, **ground soybean hulls**, calcium carbonate, dried beet pulp, salt, soybean oil, DL-methionine, pyridoxine hydrochloride, choline chloride, menadione dimethyl-pyrimidinol bisulfite (source of vitamin K), thiamine mononitrate, chole-calciferol, dicalcium phosphate, silicon dioxide, vitamin A acetate, folic acid, biotin, dl-alpha tocopheryl acetate (form of vitamin E), calcium pantothenate, riboflavin, nicotinic acid, vitamin B12 supplement, manganous oxide, zinc oxide, ferrous carbonate, copper sulfate, zinc sulfate, calcium iodate, cobalt carbonate | Ground wheat, ground corn, corn gluten meal, wheat middlings, soybean oil, calcium carbonate, dicalcium phosphate, brewers dried yeast, L-lysine, iodized salt, magnesium oxide, choline chloride, DL-methionine, calcium propionate, L-tryptophan, vitamin E acetate, menadione sodium bisulfite complex (source of vitamin K activity), manganous oxide, ferrous sulfate, zinc oxide, niacin, calcium pantothenate, copper sulfate, pyridoxine hydrochloride, riboflavin, thiamin mononitrate, vitamin A acetate, calcium iodate, vitamin B12 supplement, folic acid, biotin, vitamin D3 supplement, cobalt carbonate |
| **Calories from Protein** | 29% | 23% |
| **Calories from Fat** | 13% | 22% |
| **Calories from Carbohydrates** | 58% | 55% |
