## Supplementary material for "Notch dimerization provides robustness against environmental insults and is required for vascular integrity": S2 Fig

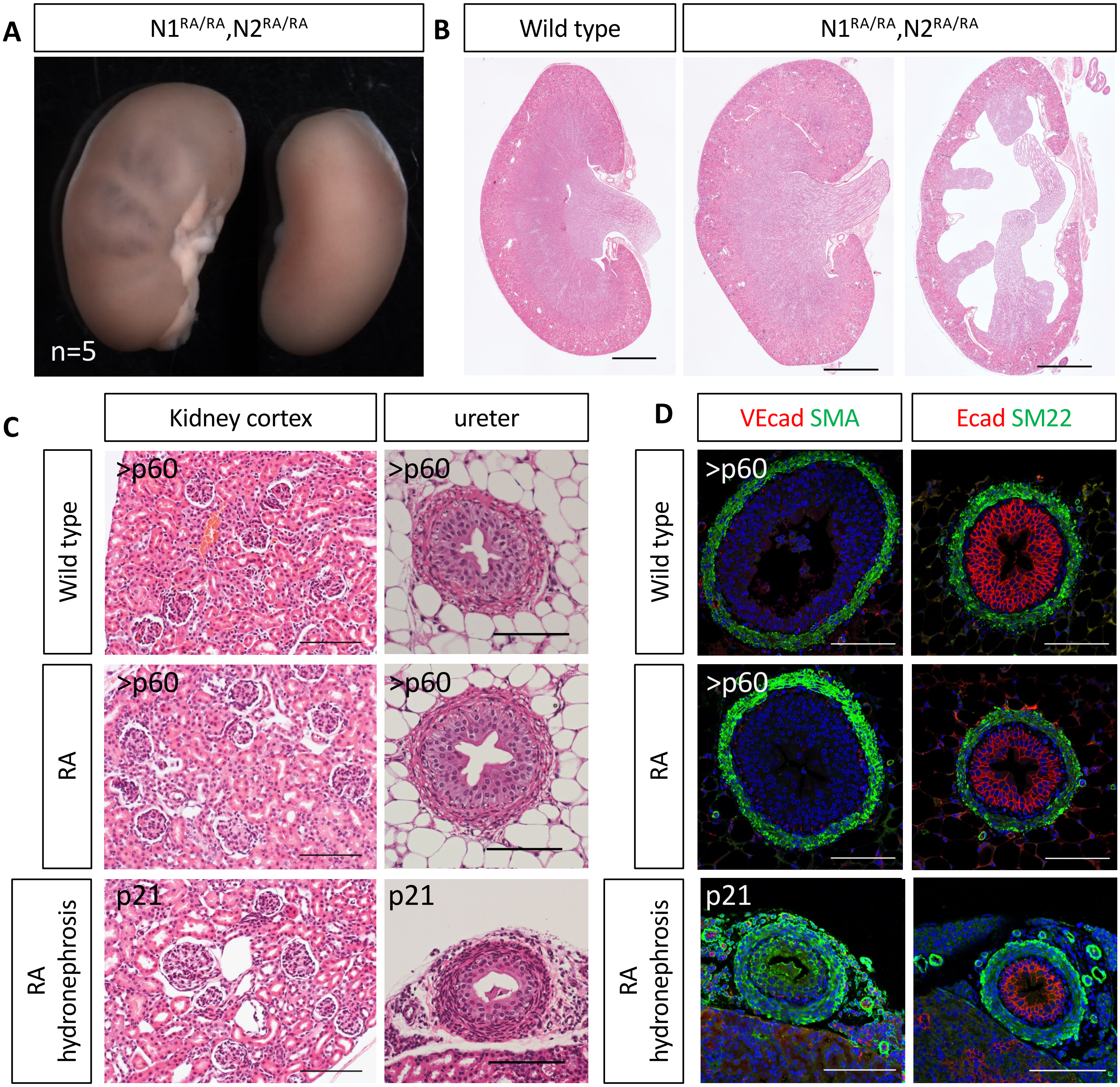


**S2 Fig. Loss of Notch dimerization can cause unilateral hydronephrosis.** (**A**) Some Notch dimer deficient mice develop unilateral hydronephrosis. (**B**) H&E stains of kidney sections from P21 WT and RA mice. Scale bars 1 mm. (**C**) Histological analysis shows no change in glomeruli density in dimer deficient mice. Scale bars 100 μm. (**D**) Ureter morphology is not altered in dimer deficient mice and the amount of smooth muscle cells around the ureter is comparable in WT and RA mice. Scale bars 100 μm.
