## Supplementary material for "Notch dimerization provides robustness against environmental insults and is required for vascular integrity": S2 Table

**S2 Table:** Chi-square test for ratio of N1$^{RA/-}$; N2$^{RA/-}$ mice (p=0.00001162) and of N1$^{+/-}$; N2$^{+/-}$ controls (p=0.7358) at birth, as well as Chi-square test comparing N1$^{RA/-}$; N2$^{RA/-}$ mice born on extruded compared to autoclaved diet (data from Kobia et al. 2020) (p=$10^{-11}$).

| **A** genotype | **expected** | | **observed** | | |
|---|---|---|---|---|---|
| | **%** | **#** | **%** | **#** | **Chi²** |
| N1:+/RA, N2:+/RA | 25 | 63.5 | 32.68 | 83 | 5.9882 |
| N1:RA/^, N2:+/RA | 25 | 63.5 | 21.26 | 54 | 1.4213 |
| N1:+/RA, N2:RA/lacZ | 25 | 63.5 | 32.28 | 82 | 5.3898 |
| N1:RA/^, N2:RA/lacZ | 25 | 63.5 | 13.78 | 35 | 12.7913 |
| **sum** | 100 | 254 | 100 | 254 | 25.5906 |

| **B** genotype | **expected** | | **observed** | | |
|---|---|---|---|---|---|
| | **%** | **#** | **%** | **#** | **Chi²** |
| N1:+/+, N2:+/+ | 25 | 25.75 | 24.27 | 25 | 0.0218 |
| N1:+/^, N2:+/+ | 25 | 25.75 | 21.36 | 22 | 0.5461 |
| N1:+/+, N2:+/lacZ | 25 | 25.75 | 29.13 | 30 | 0.7015 |
| N1:+/^, N2:+/lacZ | 25 | 25.75 | 25.24 | 26 | 0.0024 |
| **sum** | 100 | 103 | 100 | 103 | 1.2718 |

| **C** genotype | **Autoclaved diet (Kobia et al. 2020)** | | **Extruded diet (this study)** | | |
|---|---|---|---|---|---|
| | **%** | **#** | **%** | **#** | **Chi²** |
| N1:+/RA, N2:+/RA | 41.05 | 78 | 32.68 | 83 | 0.3205 |
| N1:RA/^, N2:+/RA | 24.74 | 47 | 21.26 | 54 | 1.0426 |
| N1:+/RA, N2:RA/lacZ | 26.84 | 51 | 32.28 | 82 | 18.8431 |
| N1:RA/^, N2:RA/lacZ | 7.37 | 14 | 13.78 | 35 | 31.5000 |
| **sum** | 100 | 190 | 100 | 254 | 51.7062 |
